# Nutrient gradients mediate creation of highly resistant layers in structured microbial populations during antibiotic exposures

**DOI:** 10.1101/2022.02.02.478895

**Authors:** Mirjana Stevanovic, Thomas Boukéké-Lesplulier, Lukas Hupe, Jeff Hasty, Philip Bittihn, Daniel Schultz

**Affiliations:** Department of Microbiology & Immunology, Dartmouth – Geisel School of Medicine, Hanover, NH, USA; Max Planck Institute for Dynamics and Self-Organization, Göttingen, Germany; École Normale Supérieure de Lyon, Université Claude Bernard Lyon 1, Université de Lyon, Lyon, France; BioCircuits Institute, University of California, San Diego, La Jolla, CA, USA; Department of Bioengineering, University of California, San Diego, La Jolla, CA, USA; Molecular Biology Section, Division of Biological Sciences, University of California, San Diego, La Jolla, CA, USA

**Keywords:** antibiotic resistance, biofilms, cell responses, gene regulation, dynamics

## Abstract

Antibiotic treatments often fail to eliminate bacterial populations due to heterogeneity in how individual cells respond to the drug. In structured bacterial populations such as biofilms, bacterial metabolism and environmental transport processes lead to an emergent phenotypic structure and self-generated nutrient gradients towards the interior of the colony, which can affect cell growth, gene expression and susceptibility to the drug. Even in single cells, survival depends on a dynamic interplay between the drug’s action and the expression of resistance genes. How expression of resistance is coordinated across populations in the presence of such spatiotemporal environmental coupling remains elusive. Using a custom microfluidic device, we observe the response of spatially extended microcolonies of tetracycline-resistant *E. coli* to precisely defined dynamic drug regimens. We find an intricate interplay between drug-induced changes in cell growth and growth-dependent expression of resistance genes, resulting in the redistribution of nutrients and the reorganization of growth patterns. This dynamic environmental feedback affects the regulation of drug resistance differently across the colony, generating dynamic phenotypic structures that maintain colony growth during exposure to high drug concentrations and increase population-level resistance to subsequent exposures. A mathematical model linking metabolism and the regulation of gene expression is able to capture the main features of spatiotemporal colony dynamics. Uncovering the fundamental principles that govern collective mechanisms of antibiotic resistance in spatially extended populations will allow the design of optimal drug regimens to counteract them.

## 1 Introduction

Microbial communities in their natural environments are remarkably dynamic and heterogeneous. Cellular responses and other physical and biological processes that shape these communities take place in structured environments that change over time, with the coexistence of diverse phenotypes and complex behaviors at the population level (Nadezhdin et al. 2020; Liu et al. 2015; Fu et al. 2018; Mavridou et al. 2018; Mukherjee and Bassler 2019; S. R. Scott et al. 2017; Bittihn et al. 2020). In biofilms, the predominant form of bacteria in the wild, communities are spatially structured, with high cell density and constrained mobility (Nadell, Xavier, and Foster 2009). In such dense populations, single cells interact with their immediate environment and collectively modify it, creating structures with diversified phenotypes at larger scales across the biofilm (Cao et al. 2016; Besharova et al. 2016). The coupling between bacterial metabolism and environmental factors, such as the diffusion of metabolites and other chemical compounds throughout the microcolony, alter the local availability of these compounds and create large-scale chemical gradients (Stewart and Franklin 2008). This uneven distribution of resources can affect cell growth, gene expression and susceptibility to antibiotics in different parts of the colony, leading to the emergence of phenotypic structures across the population.

One of the main advantages for bacteria associated with the biofilm lifestyle is resistance against harmful chemical compounds, such as antibiotics (Bottery, Pitchford, and Friman 2021). Biofilms are notoriously hard to treat in the clinic, commonly surviving much higher drug doses than communities of planktonic cells (Orazi and O’Toole 2019). Much of the phenotypic diversity in such structured environments results from the decreasing availability of resources that are actively consumed towards the interior of the colony (Kowalski et al. 2020). Cells closer to the surface have unrestricted access to nutrients, while cells in the interior of the colony have to adapt their metabolism to lower availability of nutrients. This phenotypic diversity is advantageous for microbial communities exposed to environmental stresses, providing alternative collective means to deal with changing conditions (Ackermann 2015; Hesper Rego, Audette, and Rubin 2017; Nguyen, Lara-Gutiérrez, and Stocker 2020; Nguyen et al. 2021; D’Souza et al. 2021; Lee et al. 2019). In the event of sudden exposure to insults, the colony can redistribute and use resources more efficiently during the activation of cell responses and adjust the spatial distribution of phenotypes to increase chances of survival (Dewachter, Fauvart, and Michiels 2019; Sánchez-Romero and Casadesús 2014). To explain the large-scale reorganization of microbial populations in variable environments resulting from the concerted interaction of diverse phenotypes, we need a quantitative understanding of how environmental cues are differently sensed and processed by single cells and integrated into complex behaviors at the community level. However, despite remarkable advances in understanding the regulation of cell responses at the molecular level, we still lack a framework to analyze how regulatory circuits control microbial behavior at the population level.

Here, we investigate how phenotypic diversity originating from environmental variations increases resistance at the population level during drug responses. We focus our analysis on the tet operon, which provides resistance against tetracycline (a translation inhibitor) in *Escherichia coli*, to understand the role of regulation on reshaping phenotypic structure upon an abrupt increase in drug concentration (Meier, Wray, and Hillen 1988; Fernández and Hancock 2012; Le et al. 2005). The expression of a tetracycline-specific efflux pump TetA is tightly repressed by the transcription factor TetR, which also represses itself (Fig. 1A). In the presence of tetracycline, TetR binds the drug and greatly diminishes the affinity for its operators, releasing expression of TetA, which then actively transports tetracycline out of the cytoplasm. The tet operon therefore provides an ideal system to study dynamic regulation; it is tightly repressed, well characterized, and controls a highly dynamic response which is crucial for cell survival upon antibiotic exposure (Schultz, Palmer, and Kishony 2017; Muthukrishnan et al. 2012; Le et al. 2006).

**Figure 1.**
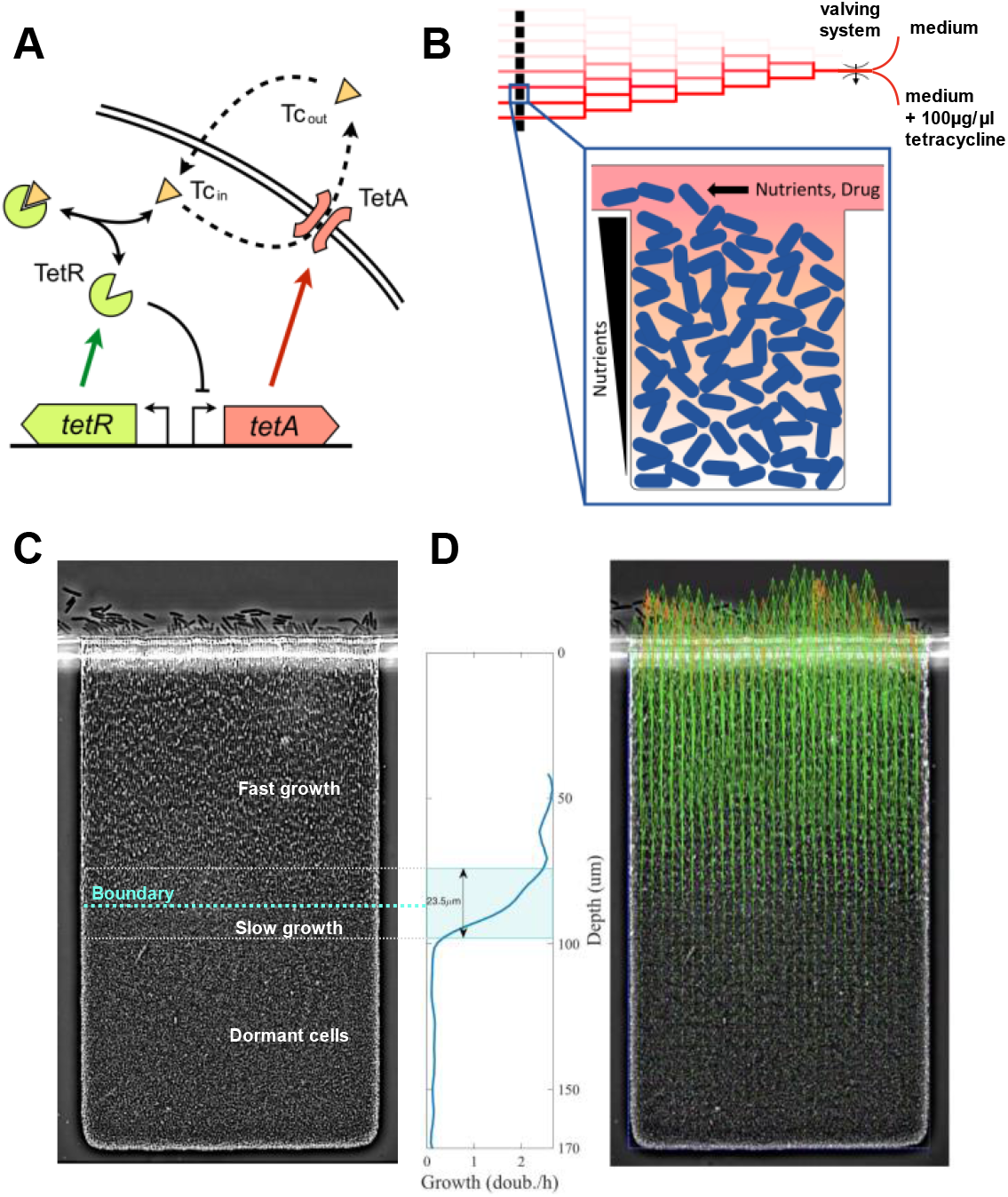
Nutrient depletion in microcolonies creates a boundary between fast-growing and arrested cells. (A) *tet* resistance mechanism: tetracycline (Tc), a translation inhibitor, diffuses across the cell membrane and binds transcription repressor TetR, which becomes inactive and releases expression of both TetR and TetA. Efflux pump TetA exports tetracycline out of the cell in an ion exchange. (B) Schematic of our microfluidic device. Media supply channels deliver nutrients to one edge of microcolonies. Nutrients diffuse down the trap and are actively consumed, thereby creating a gradient towards the interior of the colony. A linear mixer creates 8 different drug concentrations, which are quickly delivered to cell traps through the supply channels. (C) Steady state of an *E. coli* colony in the absence of drugs. Nutrient depletion creates a relatively sharp boundary that divides growing cells closer to the surface of the colony and dormant cells in the interior of the trap. Around the boundary there is a transition zone of slow growth. (D) Cell growth in the lower layers of the trap pushes cells toward the surface of the colony. We obtain the velocities of this movement at each point in the trap from time-lapse microscopy images using particle image velocimetry (PIV). Green arrows represent velocity measurements, outliers are replaced by extrapolated values (orange arrows). We then calculate cell growth by differentiating vertical velocities with respect to depth. The transition zone of 23.5μm going from 90% to 10% of maximum growth is highlighted.

## 2 Results

To characterize how expression of resistance is coordinated across structured populations during antibiotic responses, we built a microfluidic device where spatially extended, biofilm-like microcolonies are exposed to precisely defined dynamic drug regimens (Bittihn et al. 2020). The device creates confined environments for the growth of microcolonies by placing traps 170μm deep, 100μm wide, and 1.65μm thick, which can house a 2-dimensional layer of densely packed cells. These traps are connected to nutrient supply channels that continuously deliver fresh medium to one side of the growing colony while washing away spillover cells (Fig. 1B). The design of our microfluidic device allows fast switching between different media, which permits sudden exposure of the colony to a high dose of drug. When loaded into the device, *Escherichia coli* colonies grow to fill the traps. As the population receives nutrients only from one edge, cells closer to this edge grow fast, while cells towards the interior of the microcolony grow less due to decreased nutrient availability. While we do not measure nutrient availability directly, this transition from growth into dormancy was shown to be a result of nutrient gradients (Bittihn et al. 2020, see Methods). In these experiments, cells deprived of nutrients at the rear of the traps switched to a non-dividing phenotype resembling the small, round morphology in stationary-phase batch cultures (Nyström 2004), also indicating that nutrient supply is inherently diffusion limited (Fig. 1C).

We determine the phenotypic structure across a microcolony of *E. coli* cells carrying the native tetracycline resistance *tet* operon during drug responses by measuring cell growth and expression of resistance genes in real time. We measured expression of resistance using a two-color reporter plasmid indicating TetR and TetA expression (P_R_-GFPmut3 and P_A_-mCherry, (Schultz, Palmer, and Kishony 2017)). Both the resistance proteins and the fluorescence proteins are stable and not actively degraded (Reuter et al. 2020; Sophie et al. 2019), so their intracellular concentrations are determined by expression and dilution due to cell growth (we do not observe strong photobleaching, which would cause underestimation of resistance proteins levels). Therefore, fluorescence signals measure intracellular accumulation of resistance proteins, rather than their expression rates. We determined cell growth throughout the colony using particle image velocimetry (Thielicke and Sonntag 2021), a numerical method that uses correlations between images in a time series to calculate velocities in a fluid environment (Fig. 1D). Since the traps are of rectangular shape, cells only move in one direction towards the opening (which we will call the “top” of the trap, corresponding to the surface of the colony), with negligible sideways movement. Therefore, we determine the cell growth rate at each depth from the top of the trap by calculating the derivative of velocities along the vertical axis of movement. Since all relevant transport processes also vary only in one dimension along the same axis, gene expression also does not vary sideways and can be expressed solely as a function of depth. We repeated this analysis in two identical traps under the same conditions, as repeat experiments, as well as an identical trap within the same device that is not exposed to the drug, as a negative control. We tested the robustness of our measurements both by comparing different vertical sections of the trap within the same image and by comparing subsequent images of the trap during steady states both with and without the drug (Supplementary Fig. 6).

The steady-state colony structure in the absence of tetracycline features a relatively sharp boundary between rod-shaped growing cells closer to the surface of the colony and small arrested cells towards the interior. Nutrients are typically actively imported and consumed by growing cells (Jeckelmann and Erni 2020), being constantly removed from the surrounding medium. This process creates a gradient of these molecules, decreasing from the surface of the colony into the interior (Bittihn et al. 2020; Flemming et al. 2016). Given the fast timescales of the diffusion of small molecules through the lengths of the traps (in the order of 20 seconds, see Methods), nutrient diffusion and consumption are mostly in a quasi-steady state, as other relevant timescales such as regulation of cell responses or metabolic shifts are at least an order of magnitude slower. In the absence of drug, we find cells grow at similar rates across the top part of the trap, while cells in the bottom part are dormant. These regions are separated by a sharp transition zone of approximately 25 μm, which indicates sudden depletion of a limiting nutrient. We refer to this zone as the “boundary layer”. The depth of this boundary from the surface of the cell depends on the rate of consumption of the limiting nutrient by growing cells, which in turn depends on their growth rate. Therefore, we use the depth of the boundary between growing and dormant cells, defined by half-maximal growth, as a measure of the overall metabolic state of the colony (Fig. 1C). A fast-growing colony depletes resources closer to the surface, while a slow-growing colony will consume less resources and set the boundary at greater depths.

### 2.1 Exposure to tetracycline reshapes the phenotypic structure of the microcolony

Tracking changes in patterns of cell growth and expression of resistance genes across the colony during an abrupt exposure to 100μg/ml of tetracycline (which is lethal to planktonic populations), we find an initial reduction of growth among fast-growing cells near the opening of the trap, presumably causing a redistribution of nutrients towards the interior of the colony that reactivates previously dormant cells (Fig. 2). Tetracycline is a translation inhibitor, which binds ribosomes and reduces cell growth (Chopra and Roberts 2001). Therefore, immediately after exposure, cells at the surface of the colony have their growth rate significantly reduced (Fig. 3), which also decreases consumption of nutrients.

**Figure 2.**
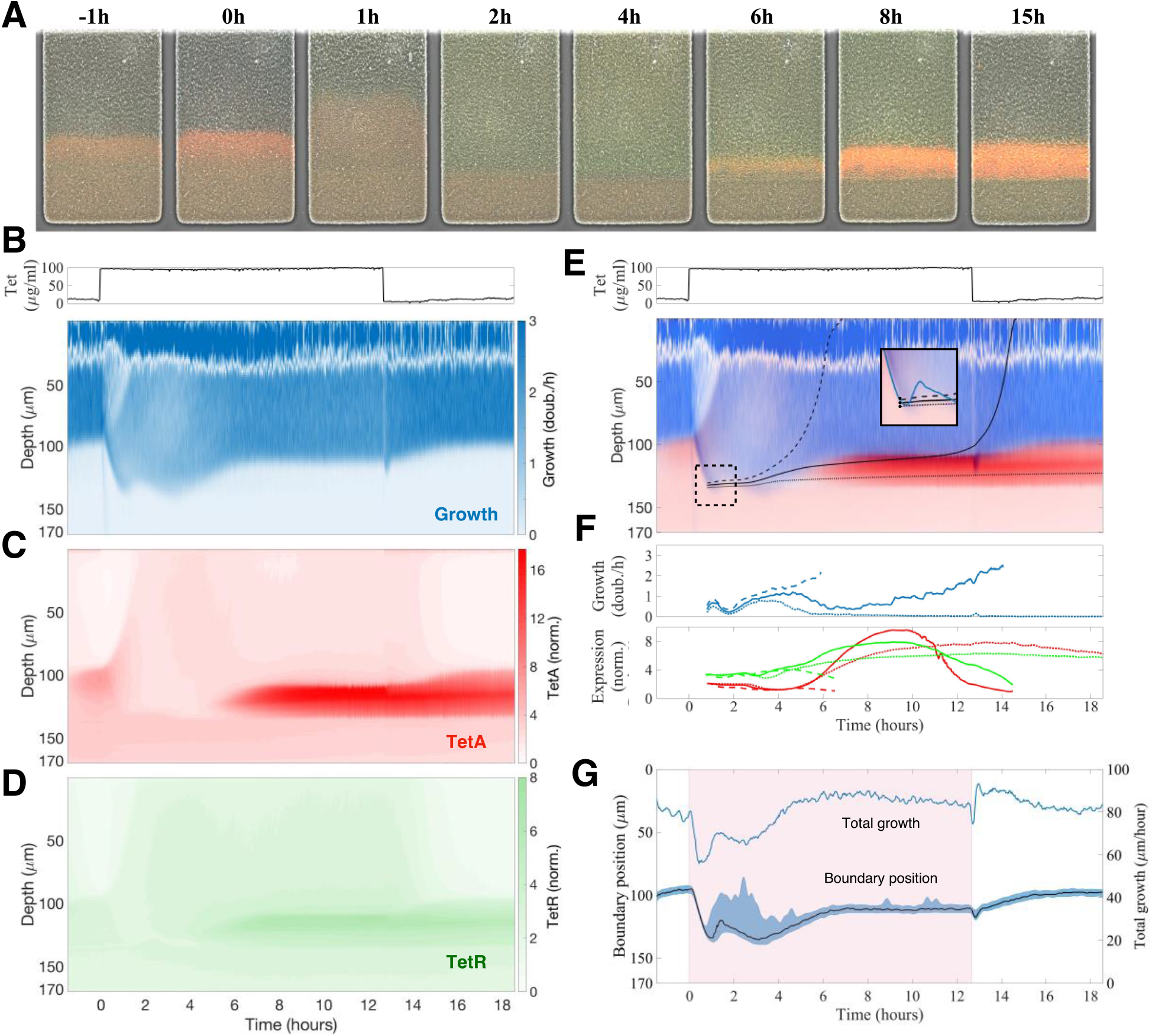
Sudden exposure to tetracycline causes a reorganization of growth and expression patterns across the microcolony. (A) Images of the colony during a response to sudden exposure to tetracycline at time zero, with expression of repressor TetR shown in green and efflux pump TetA shown in red. Following exposure, growth arrest of cells near the surface makes nutrients available deeper into the trap, so the boundary between growing and arrested cells becomes diffuse. (B-D) Kymographs of (B) cell growth, (C) efflux pump TetA and (D) repressor TetR expression during a tetracycline response, a graphical representation of spatial position over time. Since neither cell growth nor gene expression vary substantially across the horizontal dimension of the trap, we calculate the average value of each measurement at each depth, at each timepoint. These average values are denoted by the color gradients in the kymographs. Cell growth is noisy near the top of the colony due to the difficulty in calculating velocities. Expression levels of TetA and TetR are normalized to basal expression levels, obtained from fast-growing cells in the absence of drug. Above, tetracycline concentration in the medium throughout the experiment, showing the period of exposure. (E) *Single cell trajectories*: Superimposed kymograph of cell growth (blue) and TetA expression (red), with black lines showing three approximate trajectories followed by single cells, calculated from the velocities. *Inset:* these cells are initially located near each other, by the boundary, shortly after the time of exposure, but later follow divergent paths. (F) Cell growth and expression of resistance genes TetR (green) and TetA (red) along these trajectories. (G) Boundary position follows the total growth of the colony. The period of exposure to tetracycline depicted in pink shading. Position of the boundary between growing and arrested cells during the drug response, defined as the depth of half-maximal growth at each timepoint (black line). The blue shading denotes the interval between 30% and 70% maximal growth at each time point. The total growth of the colony, defined as the integral of the growth across the whole depth of the trap, is simply the velocity of cells at the top of the trap (blue line).

**Figure 3.**
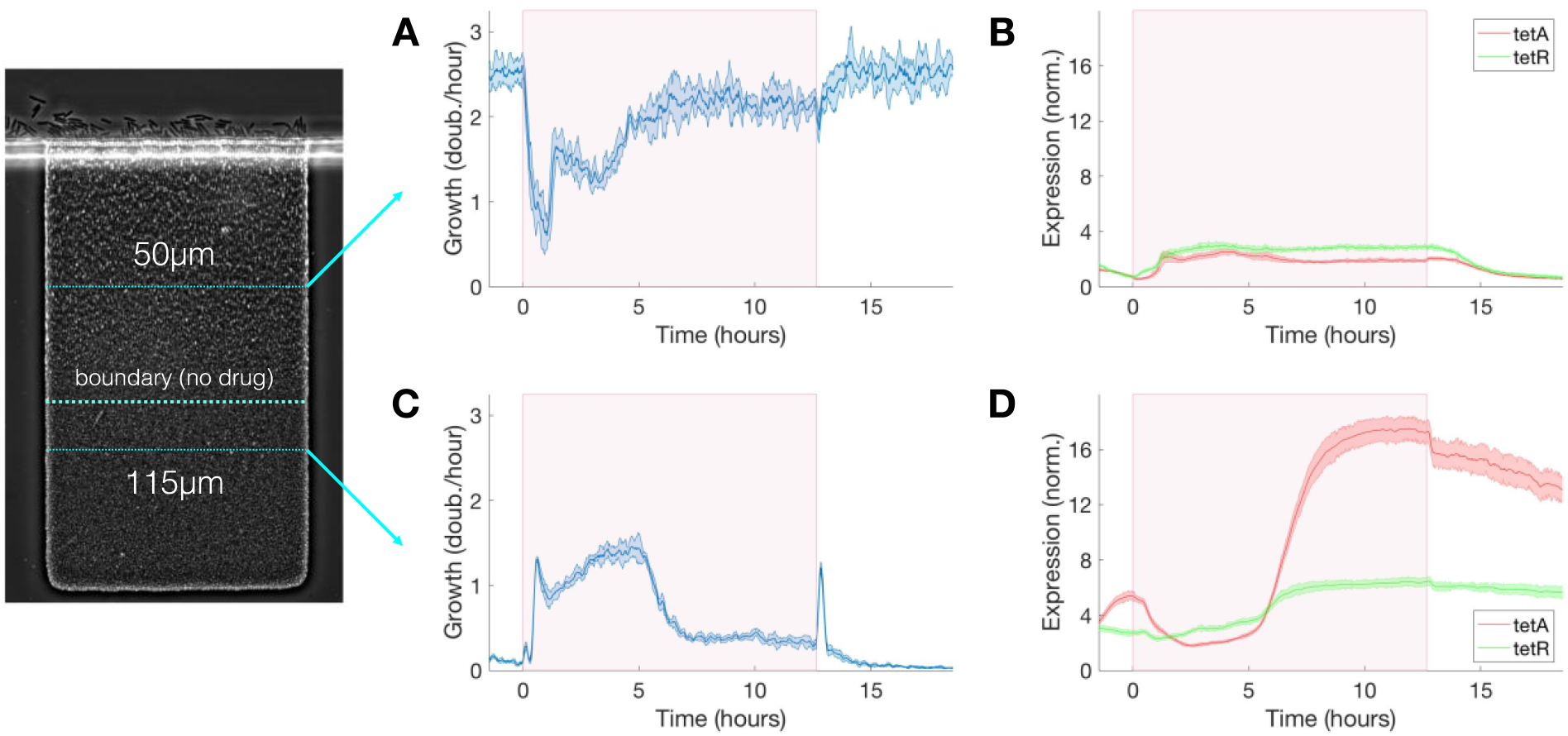
Transient growth in the interior of the colony during drug exposure induces high levels of resistance. (A) Cell growth, (B) TetA and TetR expression at a fixed depth of 50μm during exposure to tetracycline. Cell growth decreases upon exposure but is mostly recovered after 5 hours. TetA and TetR expression increase during exposure but return to pre-exposure levels after drug is removed. (C) Cell growth, (D) TetA and TetR expression at a fixed depth of 115μm during exposure to tetracycline. Transient increase in cell growth at lower depths corresponds to the period of reduced growth at the top of the trap. TetA and TetR expression increase substantially during exposure, and high expression levels are kept even after removal of the drug, when these cells become dormant. The period of exposure to tetracycline is depicted in pink shading. Shaded areas around the curves denote one standard deviation from the mean. The period of exposure to tetracycline depicted in pink shading.

Following the exposure, a rearrangement of the growth profile creates a zone of intermediate growth rates that extends deeper into the colony (Fig. 2) and temporarily reactivates cells beyond the original boundary, which were dormant before exposure. This observation is consistent with the reduced nutrient consumption at the colony surface, where the resulting surplus of nutrients becomes available to cells further inside the colony, blurring the boundary between growing and arrested cells and causing the reactivation.

After a transient period, growth at the colony surface is restored, and expression of resistance is resumed among growing cells at the colony surface. Unlike nutrients, active import of antibiotics is at least very rare (Delcour 2009). In our case, tetracycline diffuses slowly through the cell membrane, with a half-equilibration time of approximately 45 minutes (Reuter et al. 2020). Tetracycline is also not actively degraded intracellularly by enzymes, so cells do not act as a significant sink for the drug, as they do for nutrients. Since diffusion of tetracycline through the extracellular space of the trap is much faster than uptake by cells and removal by advection (diffusion time of ∼20 seconds like for nutrients, see Methods), the drug can quickly diffuse from the opening of the trap into the interior.

We confirm the timescale and reach of diffusion of tetracycline through the trap by measuring the diffusion of a dye of similar size (sulforhodamine, MW=558) added to the medium containing the drug (Supplementary Fig. 2). Therefore, upon exposure to antibiotics, all growing cells across the colony can sense the presence of the drug and induce drug responses. As TetA is expressed and growth at the surface of the colony resumes, so does the consumption of nutrients, which causes the boundary between growing and arrested cells to move closer to the opening of the trap once again (Fig. 2G). Around 5 hours after exposure, after expression of resistance and cell metabolism have stabilized to the new steady state, growth at the surface of the colony settles at a rate 75% of that in the absence of drug, and a sharp boundary is again delineated 20μm further into the trap from the location of the original boundary before drug exposure (Fig. 2BG). Upon removal of tetracycline, the growth profile of the colony returns to the same levels as before exposure within 3 hours.

The transient period following exposure to tetracycline is dominated by two processes: within the first hour, the growth profile of the colony is determined by levels of TetA already present at the time of exposure; then, in the following 4 hours, the growth profile is determined by activation of resistance in growing cells throughout the colony. Slow-growing cells around the boundary generally show higher levels of TetA expression prior to exposure (Fig. 2C). Higher levels of expression of resistance in these cells can be expected from the proteome partition, since slow-growing cells do not need to devote as many resources to the production of ribosomes (Scott et al. 2010, further discussed below in our model). Due to this growth-dependance in the expression of resistance, when fast-growing low-TetA cells at the surface of the colony are suddenly arrested by contact with tetracycline, slow-growing high-TetA cells at the boundary are invigorated with a ready supply of nutrients and increase growth to occupy the top of the colony, displacing the arrested cells (Fig. 2BC). Although preexisting TetA levels are initially reduced in these cells as they begin to grow faster, this reduction is compensated by the strong expression of TetA induced upon exposure, reaching high levels even in fast-growing cells as the colony approaches steady state (Fig. 3). Throughout this process the levels of repressor TetR change in similar fashion as TetA, reflecting their common regulation by TetR itself (Schultz et al. 2017), although fully expressed TetR does not reach the same increase in relation to basal levels as TetA does (Fig. 3).

As expression of resistance recovers growth at the top of the colony, the boundary between growing and arrested cells moves back towards the surface. The temporary activation of dormant cells deep into the colony, which happens in the transient period following exposure, induces high levels of resistance in cells growing slowly in the presence of tetracycline (Fig. 3). These cells once again stop growing when growth is restored at the surface, providing the colony with a fresh layer of dormant cells with high basal levels of resistance (Fig. 2B-D). Due to their low metabolism, these cells keep high TetA levels even after the drug is removed, suggesting a mechanism where highly resistant dormant cells can be quickly activated in the event of future drug exposures (we explore this hypothesis in the next section). The reorganization of growth patterns upon drug exposure also ensures that the colony is able to keep consuming the available resources and maintain overall growth. Despite the reduction of growth at the surface of the colony, the population as a whole experiences only a mild reduction in growth during drug exposure (Fig. 2G) as active growth is merely shifted from the surface to the interior of the colony upon drug exposure (Fig. 3).

The complex dynamics of growth and expression of resistance following drug exposure, particularly around the boundary between growing and arresting cells, can lead neighboring cells to different phenotypes. Although it is difficult to track single cells throughout the experiment, we use the local velocities at each timepoint to estimate the spatial trajectories followed by single cells during drug exposure. We follow three “single-cell” trajectories that initiate at close locations in space, near the deepest point reached by the boundary, shortly after exposure (Fig. 2EF). A first cell, whose trajectory initiates slightly above the boundary, is quickly pushed towards the surface of the colony by the transient growth of deeper layers following exposure, exiting the trap before any significant expression of TetA. A second cell, whose trajectory initiates at the boundary, is only slightly dislocated by the transient growth of deeper layers, which lasts until growth at the surface of the colony is restored. This trajectory then goes through a period of slow growth when expression of TetA reaches high levels. Eventually, this cell enters the fast-growth region above the boundary layer, reducing TetA levels on its way out of the trap. A third cell, starting right below the boundary, goes through a period of transient growth itself, when expression of TetA is high, but does not have enough push from growth of deeper layers to move significantly up the trap. This cell eventually goes back to dormancy, keeping high levels of TetA. In general, from the top to the bottom of the colony, we see the same three phenotypes observed previously in single-cell studies of drug responses: fast-growing cells expressing moderate levels of efflux pump TetA close to the surface of the colony, slow-growing cells overexpressing TetA around the boundary, and dormant cells with little TetA expression in the deep interior (Schultz, Palmer, and Kishony 2017).

### 2.2 Transient growth in colony interior increases resistance for subsequent drug exposures

The temporary reorganization of growth patterns following drug exposure suggests a colony-wide mechanism of resistance by which transient growth promotes high expression of resistance in dormant cells in the interior of the colony. These resistant cells are then able to grow and repopulate the colony during subsequent drug exposures, when fast-growing cells at the surface of the colony are arrested by sudden contact with the drug. Since the rate of expression of resistance genes depends on the cell’s metabolic state, fast-growing cells close to the surface are better positioned to regulate resistance levels to optimize growth, whether in the presence or absence of drug. However, fast-growing cells are particularly affected upon sudden drug exposures when their resistance levels are low. Therefore, dormant cells in the interior of the colony, which are not able to change their protein levels and keep high levels of resistance even in the absence of drug, can quickly start growing and replace the arrested cells at the top. In this manner, the colony can collectively maintain cell growth during drastic changes in the environment.

To study the dynamics of antibiotic responses in fluctuating environments, we subjected the colony to windows of 5 min, 10 min, 20 min, and 60 min of tetracycline exposure, delivered every two hours (Fig. 4). Regardless of the duration of drug exposure, we observe the same colony dynamics described above. Pulses of high drug concentrations permanently arrest growth in fast-growing cells at the top of the colony, which have low TetA levels at the time of exposure, while promoting growth further into the colony (Fig. 5). Over the next hour, the cells at the top of the trap are replaced by the progeny of cells located below the boundary at the time of exposure, which already have significant levels of TetA upon contact with the drug (Fig. 4AB). This process is initiated even by short pulses of drug and continues to take place in the absence of the drug. This suggests both that growing cells are permanently arrested shortly after contact with tetracycline and that initial recovery of colony growth does not depend on new expression of TetA, but on the presence of cells with preexisting high levels of resistance.

**Figure 4.**
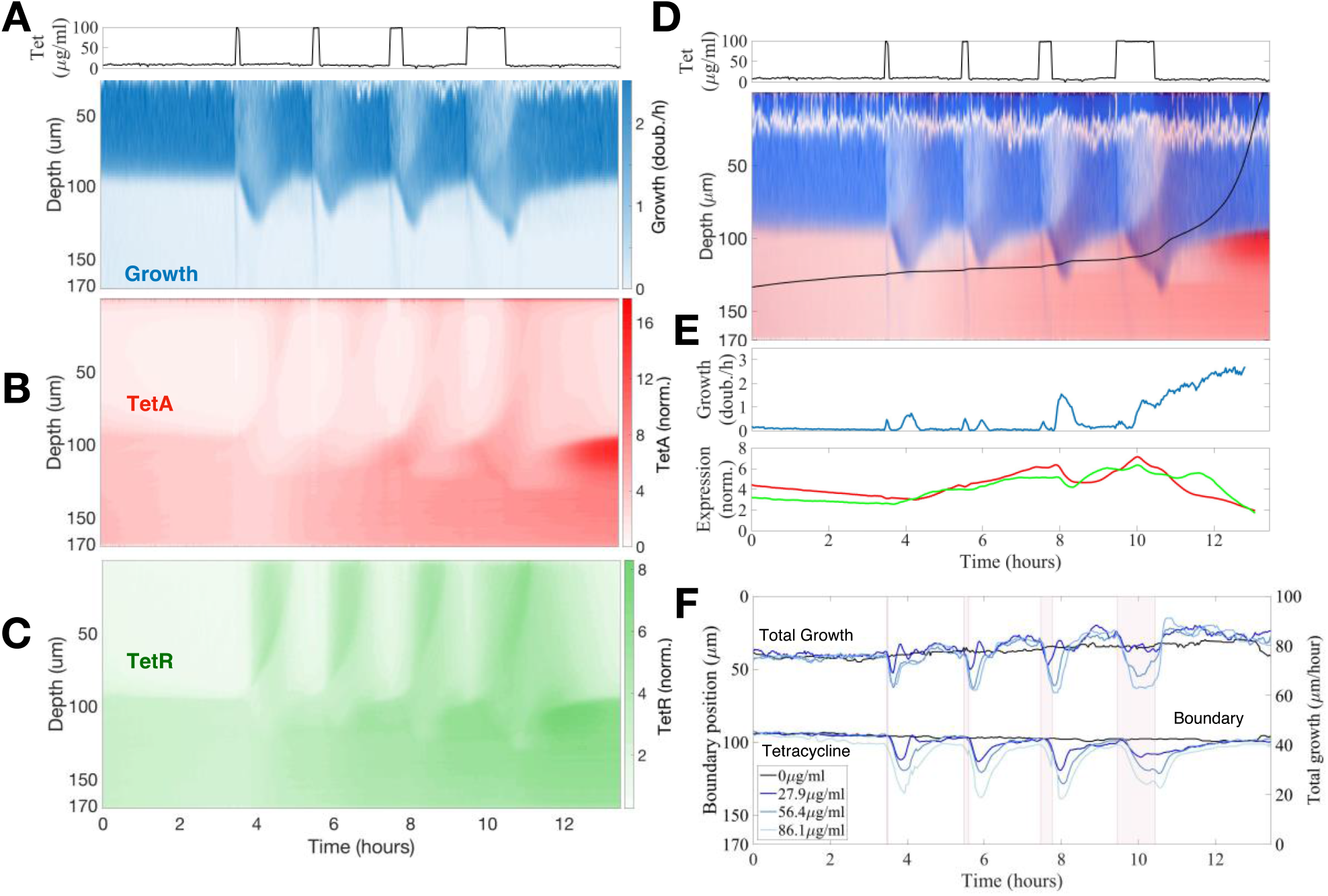
Previous drug exposures build up resistance in the colony interior to resist future exposures. (A-C) Kymographs of (A) cell growth, (B) efflux pump TetA and (C) repressor TetR expression during a sequence of tetracycline exposures, with pulses of 5 min, 10 min, 20 min and 60 min delivered every 2 hours. Expression levels of TetA and TetR are normalized to basal expression levels in fast-growing cells in the absence of drug. Above, tetracycline concentration in the medium during the experiment. (D) *Single cell trajectory*: Superimposed kymograph of cell growth (blue) and TetA expression (red), with a black line showing an approximate trajectory followed by a single cell originating in the interior of the colony, calculated from the velocities. (E) Cell growth and expression of resistance genes along this trajectory. Cell growth is noisy near the top of the colony due to the difficulty in calculating velocities. (F) Position of the boundary between growing and arrested cells during drug exposure, defined as the depth of half-maximal growth at each timepoint. Total growth of the colony during the sequence of exposures, defined as the integral of the growth across the whole depth of the trap. Periods of exposure to tetracycline are depicted in pink shading. The different lines denote exposure to different drug concentrations.

**Figure 5.**
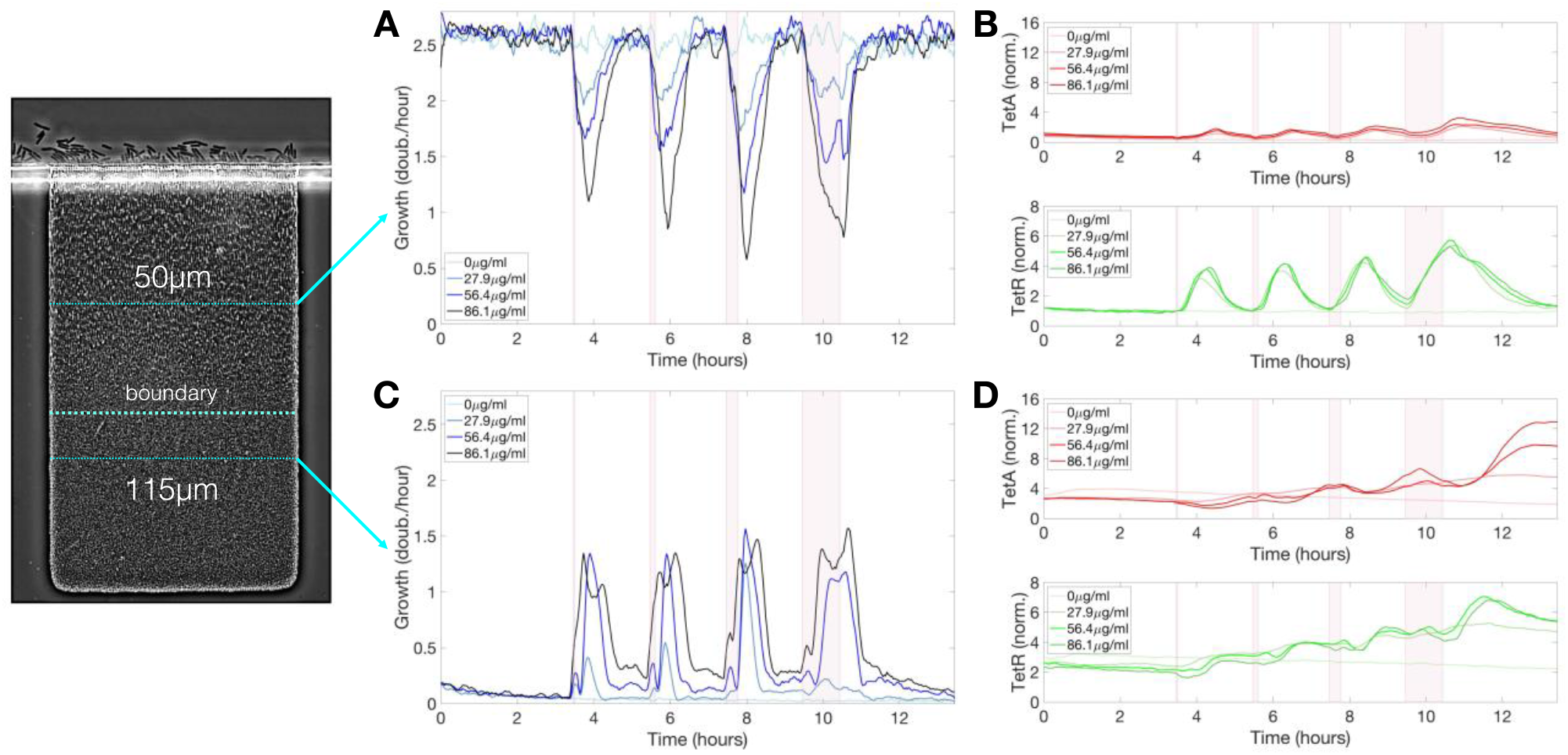
Sequence of exposures to tetracycline builds up resistance among slow-growing cells below the boundary. (A) Cell growth, (B) TetA and TetR expression at a fixed depth of 50μm during a sequence of exposures to tetracycline, with pulses of 5 min, 10 min, 20 min and 60 min delivered every 2 hours. Cell growth decreases upon each drug exposure for about 2 hours, regardless of exposure duration, but later recovers. TetA and TetR expression increase following each exposure, but later return to pre-exposure levels. (C) Cell growth, (D) TetA and TetR expression at a fixed depth of 115μm during a sequence of exposures to tetracycline. Transient increases in cell growth at lower depths correspond to the periods of reduced growth at the top of the trap. TetA and TetR levels increase further with each exposure. Expression levels are kept between exposures while these cells become dormant. The periods of exposure to tetracycline are depicted in pink shading.

Each pulse of tetracycline was followed by temporary expression of resistance genes at the top of the colony, which had mostly subsided by the time of the next pulse (Fig. 4BC). While TetR was expressed quickly and strongly regardless of pulse length, TetA concentration was higher after longer drug exposures, but concentration of either resistance gene was quickly reduced in growing cells in the absence of drug. On the other hand, each pulse of tetracycline permanently increased expression of resistance in slow-growing cells below the boundary (Fig. 5). Periods of transient growth following each exposure build up resistance in the slow-growing layer of cells below the boundary, which is not diluted in the absence of drug when these cells return to dormancy. This effect is strengthened during exposures to higher drug concentrations, which induce longer periods of transient growth following exposure, and reach dormant cells deeper into the colony (Fig. 4F). Similar transient growth dynamics were observed for short drug pulses of any length, which were sufficient to permanently arrest fast-growing cells at the surface of the colony.

We follow a single-cell trajectory initiating below the boundary at the beginning of the experiment. Slow-growing cells deep into the colony already show higher expression of resistance genes than fast-growing cells even before first contact with the drug. With each pulse of tetracycline, transient growth at the bottom of the colony moves this cell towards the surface, while increasing TetA levels (Fig. 4DE). Immediately before the last exposure, this trajectory is right below the boundary, with high TetA levels. The cell following this trajectory then resists the exposure, keeps growing, and quickly moves to occupy the top of the colony. This trajectory shows the buildup of resistance by slow-growing cells in the interior of a colony subjected to frequent exposures to the drug, which allows a quick transition into fast growth when an exposure causes nutrients to become available to them.

### 2.3 Slow-growing cells express higher levels of resistance genes

The collective dynamics of the drug response in the microcolony is characterized by markedly different gene expression levels between fast- and slow-growing cells. To determine the functional relationship between cell growth and expression of TetA and TetR, we correlated gene expression with cell growth across the trap during steady states, both in the absence and presence of tetracycline. In the absence of drug, we find that TetA and TetR expression decreases linearly with the cell growth rate (Fig. 6, Supp. Fig. 8). This linear relationship is predicted by the theory of proteome partition, whereby for a cell to grow faster, it needs to dedicate a larger portion of its protein synthesis capacity for ribosome production (up to 50%), leaving less resources for the remaining portion of its proteome (Scott et al. 2010).

**Figure 6.**
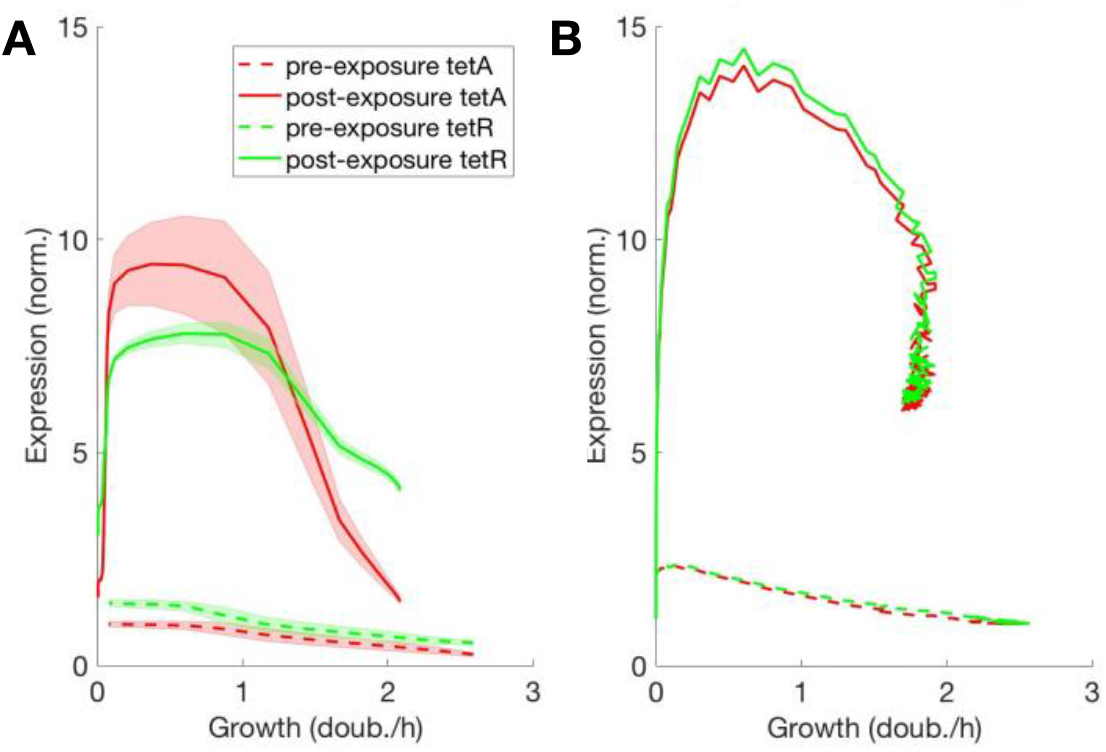
Steady-state expression of resistance decreases linearly with cell growth. Functional relationship between expression of TetA/TetR and cell growth, obtained from the steady states with and without drug, both (A) measured in the trap and (B) simulated from the mathematical model. TetA and TetR expression decrease linearly with the cell growth rate, as predicted by the theory of proteome partition. At extremely low cell growth, both TetA and TetR expression remain low even in the presence of drug.

A linear relationship between TetA/TetR and cell growth is also found among growing cells in the presence of drug, although at much higher expression levels and with a higher slope for TetA. This further increase of TetA expression in slow-growing cells, in comparison to TetR, might reflect the different steady states reached in the interplay between drug import, TetA expression and TetR repression at different growth rates (Møller et al. 2016). At lower growth rates, a decrease in dilution of intracellular drug could be compensated by increased export by TetA. However, it should be noted that high-TetA slow-growing cells around the boundary only go through a few division cycles (∼6) on their way out of the trap, and therefore gene expression is not likely to reach equilibrium at any given growth rate during this process. At extremely low cell growth in the back of the trap, both TetA and TetR expression are low even in the presence of drug, since these cells do not experience any growth and largely keep the same protein levels as before drug exposure.

### 2.4 A mathematical model linking metabolism and gene expression captures the main features of spatiotemporal colony dynamics

Both the dynamics of antibiotic responses and the resulting changes in the phenotypic structure of the colony are ultimately controlled by gene regulatory circuits (Schultz et al. 2009). However, upon environmental shifts, the progression of cellular responses does not only depend on the direct regulation provided by transcription factors but also on global effects on protein expression linked to the metabolic state of the cell (Klumpp, Zhang, and Hwa 2009; Scott et al. 2010). Failure to quickly deploy resistance genes results in higher concentrations of intracellular drug and further reduction in expression of resistance genes and drug accumulation, a positive feedback known to result in the coexistence of growing and arrested cells in the presence of antibiotics (Deris et al. 2013). Therefore, gene regulatory mechanisms control expression of resistance across the colony both directly by regulation and indirectly by the reorganization of growth patterns across the colony resulting from the expression of resistance and nutrient redistribution (Stewart et al. 2019).

To test whether coupling between direct resistance regulation, metabolism and environmental interactions is indeed sufficient to explain the complex spatiotemporal dynamics described above, we set up an agent-based computational model of growing rod-shaped bacteria incorporating these three ingredients. Each cell in the simulation is equipped with a gene regulatory circuit for the expression of resistance genes, where growth and expression are sensitive to both drug concentration and nutrient availability, and nutrient concentration in the environment is tracked separately by a continuous field throughout the trap (see Methods and Supp. Fig. 2 for details). The regulatory part of the mathematical model closely follows the previously established resistance mechanism (Schultz, Palmer, and Kishony 2017) and includes the proteome allocation mechanism (Scott et al. 2010) for growth-dependent expression (also see discussion).

In the absence of the drug, the colony reaches a steady state similar to our experimental system (Fig. 7A; first snapshot), where cells close to the outlet at the top grow (blue cells) while the cells deep in the trap are deprived of nutrients and stop growing (white cells). We then subjected our virtual colony to two pulses of drug: A short 30min pulse and a long 500min pulse. These pulses triggered changes in the growth pattern and nutrient distribution (Fig. 7ABE) as well as expression of the resistance genes TetA and TetR (Fig. 7CD).

**Figure 7.**
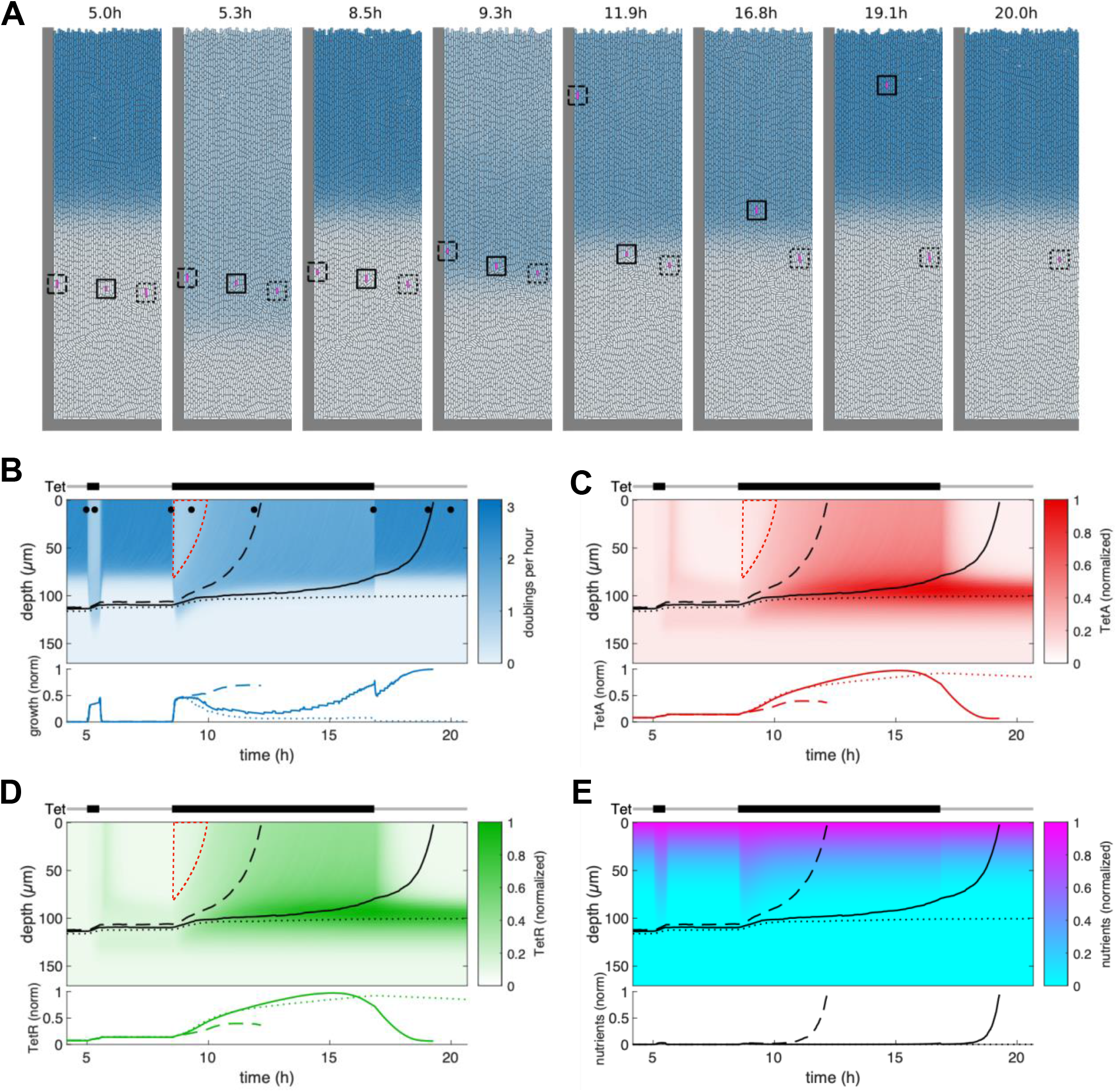
Mathematical model recovers main features of colony dynamics during drug response. (A) Snapshots of agent-based simulations, showing transient reorganization by the growth pattern during a short drug pulse (t=5.3h) and during a long pulse (t=9.3h) eventually reaching a new drug-induced steady state (t=11.9h), before returning to the original steady state after drug removal (t=19.1h). Blue color indicates growth rate. Cells tracked to yield the time traces in panels B-E are enclosed by a square of the same line style. (B) Kymograph of the growth rate over the course of the simulation. Tet exposure is indicated above. Dots at the top of the kymograph indicate time points of the snapshots in panel A. Lines indicate trajectories of three tracked cells (cf. panel A), time traces of their growth rate are shown below the kymograph. Dashed red line marks the cells in the top (previously growing) part of the cell trap which are subsequently replaced by reactivated dormant cells already primed with TetA and TetR. (C-E) Kymographs and single-cell traces as in panel B for TetA, TetR and nutrients. The full spatiotemporal dynamics can be seen in Supplementary Movie 2.

Upon exposure of the colony to the shorter first pulse, we see two main effects: First, growth is suppressed in the growing part of the population, which leads to diminished nutrient consumption and a redistribution of nutrients deeper into the trap (Fig. 7E). This causes dormant cells deeper in the trap to resume growth, albeit at a reduced rate (Fig. 7A, t=5.3h; Fig. 7B). When the pulse is over, the growth pattern returns to its original state (Fig. 7A, t=8.5h). However, the expression of resistance genes triggered by the drug induces a lasting change: When cells in the lower part of the trap return to their dormant state, their TetA and TetR concentrations are still elevated, leading to a reservoir of dormant cells with active resistance. Consequently, these cells are able to tolerate the next pulse with a higher growth rate, while growth of cells at the top suppressed to the same extent as in the naive colony (wedge-shaped region outlined in red in Fig. 7BCD).

During the longer pulse, we can see the adaptation process more clearly: The growth boundary initially deepens (albeit slightly less than during the first pulse due to the reservoir of resistant cells), but eventually settles at an intermediate position, when expression of resistance in the growing cells has reached its steady state (Fig. 7A, t=11.9h; Fig. 7B). Expression is enhanced in cells that grow slowly due to nutrient deprivation, leading to the creation of a reservoir of cells with very high levels of resistance after the steady state under drug exposure is reached (Fig. 7CD). Again, this reservoir is maintained even when the colony returns to its original growth pattern upon drug removal. The full spatiotemporal dynamics can also be observed in Supplementary Movie 2.

In contrast to the experiment, we had the ability to track individual cells throughout the simulation, since they are represented by discrete units. We chose three initial cells and extracted their growth rate, TetA and TetR concentrations as well as the nutrient availability at their respective positions (squares in Fig. 7A; Supplementary Movie 2; corresponding trajectories below each kymograph in Fig. 7B-E). The initial positions were chosen similarly to the experiment, such that the deepest cell, despite moving toward the top during transient growth, stays in the dormant part of the population, while the middle cell is expelled from the colony after drug removal, and the top cell leaves the colony even earlier, shortly after drug exposure during the long pulse. The time traces clearly show transient growth for the bottom cell and permanent reactivation of growth for the top cell upon exposure (bottom of Fig. 7B). In contrast, the middle cell experiences a complex sequence of environments during the long pulse: First, growth is triggered by increased nutrient availability. While the cells in the upper growing part of the population recover, the cell is again deprived of nutrients, leading to reduced, but non-zero growth. Simultaneously, resistance expression is markedly increased (bottom of Fig. 7CD). Finally, the residual growth allows the cell to reach the region of the trap with high nutrient availability, which both increases its growth rate and leads to a decline in resistance levels until the cell leaves the trap.

## 3 Discussion

Spatial structure results in organization of the colony with different phenotypes towards the interior, with different resistance profiles. This phenotypic structure changes during dynamic responses and determines resistance at the colony level. Microcolony microfluidics is an essential tool to describe how the reorganization of chemical gradients and growth patterns across bacterial colonies coordinate antibiotic responses at the population level. In these experiments, we described how the interplay between drug action, cell growth and expression of resistance defines the course of antibiotic responses in single cells and ultimately generates the large-scale phenotypic structures that dictate colony-level resistance during drug responses (Craig MacLean et al. 2010; Kim et al. 2020).

Notably, the rearrangement of growth patterns allowed the colony to maintain growth and coordinate expression of resistance upon a sudden exposure to a concentration of drug large enough to immediately arrest the fast-growing cells at the surface of the colony. This same concentration of drug is also sufficient to arrest growth of planktonic populations in liquid cultures for up to 20 hours (Schultz, Palmer, and Kishony 2017). Since the exposure to large doses of drug can overwhelm cells before expression of resistance genes can be accomplished, a strategic reserve of slow-growing cells with high levels of resistance proteins ensures that the colony can remain active, replacing lost cells at the surface and further expressing resistance (Lin et al. 2021). With each exposure to the drug, transient growth increases resistance levels at the base of the colony. Therefore, the colony can adjust its resistance levels over time to the regimens it typically encounters (Mathis and Ackermann 2016).

Our model was able to recapitulate the qualitative dynamics of spatially heterogeneous resistance expression and growth pattern modulation—most notably the reactivation of dormant cells upon drug exposure, the adaptation process during resistance expression that leads to an altered steady state in the presence of the drug and the persistent accumulation of resistance in the dormant subpopulation. This, together with the excellent agreement of single cell traces with the experimental findings, can serve as evidence that the ingredients of our model and their mutual coupling are indeed sufficient to explain the complex dynamics observed experimentally. It is worth noting that, in addition to nutrient diffusion, resistance expression and growth modulation by the drug and through nutrients, we found it necessary to include the dynamic allocation of resources to different protein sectors suggested by (Scott et al. 2010) as an additional ingredient. Without it, the preferential expression of resistance in nutrient-deprived cells and, consequently, the efficient priming of dormant cells for future drug exposure was not present.

One aspect that is poorly captured by our model is the time scale of the transition between different metabolic states. Both the reactivation of dormant cells upon exposure as well as the establishment of a new steady state happen on a considerably longer time scale in the experiment compared to the simulations (compare Figs. 2B and 7B). However, the consequences of transitions between different metabolic states in bacteria for growth, expression and metabolism are complex. For example, the exact role and underlying physiology of bacterial lag phase alone are a topic of active research with many open questions and could be relevant here (Bertrand 2019). Therefore, it is not surprising that our simplified model does not fully capture the complexity and time scale of these transitions. In-depth experiments and a possible refinement of the model will be necessary to disentangle the contributions of general physiological effects and specific resistance dynamics during these transitions.

Biofilms can simultaneously deploy multiple strategies to survive in fluctuating environments. While fast-growing cells at the surface of the colony can respond and adapt more quickly to environmental changes, slow-growing cells in the interior can show distinct expression profiles that are adjusted over time, to guarantee survival and repopulation of the colony upon sudden exposure to harsh conditions. The presence of both phenotypes, even though in our case it is caused by physical nutrient limitation, can be viewed as a bet-hedging strategy that increases flexibility of the colony to survive a large array of conditions. In the context of antibiotic resistance, the low metabolism of cells in the interior of the colony can increase tolerance to a variety of drugs, even in the absence of a dedicated mechanism of resistance (Crabbé et al. 2019; Donnert et al. 2020).

Although the cells in our system do not present all the phenotypic traits of cells in true biofilms, such as matrix production, we believe that the collective behaviors we observe are largely generalizable to biofilms, being driven by transport phenomena and nutrient consumption. Our system could further be used to analyze alternative scenarios that could result in different collective dynamics. In particular, the use of a bactericidal drug could test the trade-off between expression of resistance in growing cells at the top of the trap and drug tolerance in dormant cells at the bottom. Many resistance mechanisms can actively degrade antibiotics, making resistant cells act as sinks that continuously remove drug from the extracellular space (Flemming et al. 2016). This scenario could generate drug gradients on top of nutrient gradients, further protecting cells in the bottom of the trap and resulting in different phenotypic structures.

True biofilms show an even richer complexity, with phenotypic structures being shaped by several bioactive compounds secreted by the cells (Dietrich et al. 2008). Nutrient gradients, particularly relating to oxygen availability, were shown to generate subpopulations that are resistant to antibiotics (Beebout et al. 2021; Kowalski et al. 2020). Our work provides a framework to understand the complex collective dynamics of structured populations that emerge from processes that are generally studied at the cellular level. Our results suggest that the emerging phenotypic structures are determined by how metabolic and transport processes interact at larger scales, while single cells moving through the population adopt the local phenotype without disrupting large-scale structures.

These experiments with growing finite populations under controlled dynamical conditions help bridge the gap between understanding the dynamics of drug responses at the single-cell level and understanding the resulting effect on the population-level behavior. A model-based, quantitative description of how microbial populations adapt the regulation of cell responses to complex dynamical environments is crucial to understand community-level behaviors such as antibiotic resistance, pathogenesis, and biofilm formation, as well as generating synthetic systems for biotechnology applications (Xie and Fussenegger 2018; Chait et al. 2017). The collective strategies of antibiotic resistance described here are important to consider when designing clinical treatments for microbial infections, since biofilms are notoriously hard to clear.

## 4 Materials and Methods

### Media, Drugs, Strains

All strains were derived from *E. coli* K-12 strain MG1655 *rph*+ Δ*lacIZYA*. The native *tet* resistance mechanism from the Tn*10* transposon was ordered from Genewiz in a pIT3-CH integrating plasmid and integrated in the chromosome at site HKO22. Matching fluorescent reporters for TetR and TetA (GFPmut3 and mCherry, respectively), were also ordered from Genewiz in a pZS1 plasmid and transformed using TSS (Schultz, Palmer, and Kishony 2017). The pZS1 vector has a pSC101 origin of replication with stringent control, which provides good accuracy in fluorescence measurements due to low copy number variation (10 to 12 copies per cell, (Lutz and Bujard 1997)). All experiments were performed in EZ-rich defined medium (Teknova) prepared according to the manufacturer’s instructions with 0.2% glucose as the carbon source. To prevent cell adhesion to microfluidic channels, all media were supplemented with 0.075% (w:v) Tween 20. Tetracycline solutions were freshly made from powder stocks (Sigma) and filter-sterilized before each experiment. All media containing tetracycline also had 0.008% (v:v) sulforhodamine dye added, so drug concentration inside the chip could be calculated from microscopy images.

### Microfluidics

To image spatially extended micro-colonies of tetracycline-resistant *E. coli*, we used a microfluidic device that consists of multiple 100μm by 170μm traps (Bittihn et al. 2020). The night before the experiment, a culture was inoculated from a −80 °C glycerol stock into lysogeny broth (Difco) supplemented with the appropriate selection antibiotics and grown overnight in a shaking incubator at 37 °C. On the day of the experiment, the saturated overnight culture was diluted 1:1,000 into 5 ml of the same medium supplemented with 0.075% (w:v) Tween 20 and incubated until an OD600 of 0.1 was reached. Then, 1 ml of this culture was spun down at 3,000g in a standard tabletop microcentrifuge for 3 min, and the cells were resuspended in 5 μl of the base medium used for the microfluidic experiment. The cell suspension was then immediately loaded into the waste port of the microfluidic device, which had been degassed in a vacuum chamber for at least 30 min, and base medium was loaded into all other ports. The experiment was performed at 37°C. After the traps had been seeded, media and waste ports were connected to the corresponding reservoirs and media channels were flushed with base medium at a high flow rate (>2,000 μm s^−1^) before calibrating the flow rate to 100 μm s^−1^ in the media channels. A linear on-chip mixer was used to mix growth media from two inputs in 8 different proportions to achieve different drug concentrations. One of the inputs can be switched in real-time between growth media containing no tetracycline and growth media containing tetracycline to expose colonies to abrupt shifts in drug concentration. Supplementary Movie 1 shows a microcolony subjected to the tetracycline exposure regimens described in Figures 2 and 3.

### Microscopy

A Nikon Eclipse Ti2 inverted microscope was used with a Photometrics CoolSNAP HQ2 CCD camera to capture images and a Lumencor SOLA SE LED Light Engine for fluorescent excitation. The stage holding the microfluidic device was enclosed in a plexiglass incubation chamber, maintaining a constant 37 °C environment. The acquisition software was NIS-Elements. High-resolution data was acquired at ×30 magnification, with exposure times of 20 ms for phase contrast, 100 ms for GFP (25% LED intensity) and 500 ms for mCherry (20% LED intensity). With one image for each trap size and medium composition (16 images), each containing two traps, an imaging interval of 2 min was possible.

### Microscopy Data Analysis

GFP and mCherry fluorescence was extracted from microscopy images using custom-written MATLAB scripts. Drug concentration at each time point was calculated by extracting the fluorescence of the medium flowing through the medium channel above the traps, as media containing tetracycline also contained a fluorescent dye. Phase contrast images were stabilized using the Image Stabilizer plugin in ImageJ. The velocity of cells was estimated from the stabilized images using the PIVlab tool in MATLAB (Thielicke and Sonntag 2021). All images were pre-processed, regions of interest (ROIs) encompassing the full traps were selected manually, and three interrogation areas were used to calculate velocity vectors. Estimates of velocity vectors were calibrated using reference distances in the microfluidic device, and the estimated vectors were validated in post-processing by restricting velocity limits. The local cell growth rate at different depths of the trap was estimated from the vertical components of velocity vectors. Assuming that cells in the microfluidic trap are densely packed and incompressible, the velocity *v*(*x, t*) at a depth *x* is determined by the instantaneous, cumulative cell growth rate in the trap 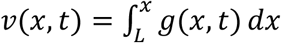. Therefore, we calculate the local growth rate by differentiating the velocity 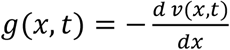.

### Estimation of the timescales of transport processes

Both nutrients and drug penetrate the trap by two processes: diffusion through the medium and advection, where inflow of medium into the trap counters the outflow of cells. This advection process can both bring nutrients and drug into the trap with the inflowing medium and remove whatever was imported by the outflowing cells. Here, we estimate the relative importance of these phenomena in determining nutrient and drug concentrations along the trap. Assuming a fast cell growth rate of 3 doublings per hour (an overestimation) and a depth of the growing layer of 100μm (as seen experimentally), the maximum advection velocity at the top of the trap is 300μm h^-1^. Therefore, the transfer time of cell mass from the growth boundary to the top of the trap is on the order of 20min, and similarly for nutrients for the counter-flow inwards for a volume fraction on the order of 0.5. The value for the diffusion constant of nutrients found to be consistent with our experimental observations for this system is D = 500μm^2^ s^-1^ (Bittihn et al. 2020), which is in line with known diffusion constants (Ma et al. 2005; Stein and Litman 1990), a recent study on biofilm front propagation (Wang, Stone, and Golestanian 2017) and the general order of magnitude for small molecules in water. Therefore, the typical time for nutrients to travel 100μm from the top edge to the growth boundary simply by diffusion is (100μm)^2^/D = 20s. Thus, an upper bound for the Péclet number in this system is Pe = 20s/20min = 1/60, which leads us to conclude that this system is diffusion-dominated and we can neglect mass transfer by advection in the growth medium. In addition, the advection velocities decrease towards zero at the growth boundary, below which only the (very fast) diffusion acts. Since diffusion constants for antibiotics are typically on the same order or even larger (tetracycline has a molecular weight of 444), this applies to all the relevant substances (nutrients and antibiotics) in our system. We also confirm the timescale of diffusion experimentally by measuring the penetration of a dye of similar size (sulforhodamine, MW=558) added to the medium containing the drug (Supplementary Fig. 2).

### Mathematical Model

We model the main biochemical interactions involved in the response as a set of differential equations that describe changes in the concentrations of the intracellular drug (*d*), the efflux pump TetA (*a*), and the repressor protein TetR (*r*):

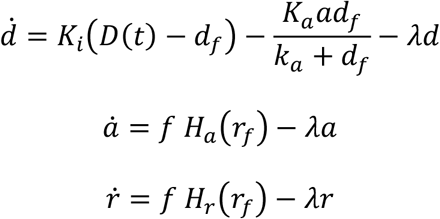

where *K*_*i*_ stands for the import rate, D for extracellular drug concentration, *d*_*f*_ for free intracellular drug, *K*_*a*_ for the catalytic rate constant of TetA, *k*_*a*_ the Michaelis constant, respectively, *r*_*f*_ free repressor, and *H*_*a*_(*r*_*f*_) and *H*_*r*_(*r*_*f*_) the synthesis rates for TetA and TetR that depend on free TetR. Since the biochemical binding and unbinding of the substrate to the transcription factor typically happens at a much faster rate than the aforementioned processes, we consider their unbound (free) forms (*d*_*f*_, *r*_*f*_) to be in chemical equilibrium with the bound form [*rd*] with a dissociation constant *K*_*d*_, such that *r*_*f*_*d*_*f*_ = [*rd*]. The term *K*_*i*_(*D*(*t*) − *d*_*f*_) represents the diffusion of drug across the membrane into the cell, 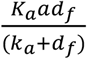 the export of drug out of the cell by the efflux pump. The terms *λd, λa* and *λr* represent the dilution due to cell growth of drug, TetA and TetR, respectively, where *λ* is the current growth rate. Both *f*, which allows us to modulate the strength of gene expression, and the growth rate *λ* depend on the nutrient level and the intracellular drug concentration.

To model these dependencies, we used the framework introduced by (Scott et al. 2010). In this framework *κ*_*n*_ and *κ*_*t*_ are the nutritional and translational capacity of the cell, respectively. Here, we refer to their base values (full nutrients, no drug) as 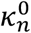 and 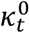, respectively.

For the nutritional capacity, we assume a Monod dependency on the limiting nutrient concentration *c* with 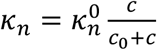, where *c*_0_ is the half-maximum concentration, which we assume to be much smaller than the nutrient concentration at the boundary of the trap in order to obtain the sharp growth boundary. For the translational capacity, we assume inhibition by tetracycline according to 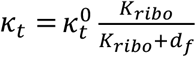, where the fraction represents the fraction of free ribosomes in the cell, and *K*_*ri*bo_ is the dissociation constant for tetracycline and the ribosome. Lower *K*_*ri*bo_ values correspond to stronger inhibition, and vice versa. According to (Scott et al. 2010), the growth rate can then be calculated as 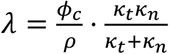, where *ϕ*_*c*_ = 0.48 and *ρ* = 0.76 are universal constants.

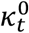 has a universal value of 4.5 *h*^−1^for *E. coli*. 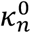 varies according to nutrient quality. We chose 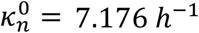 such that 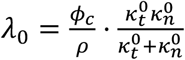 (corresponding to a state without tetracycline, at full nutrients) yields a maximum growth rate of *λ*_0_ = 0.029 *min*^−1^ as observed in the experiment (The full range of growth rates during the experiment are shown in Supp. Fig. 1).

We assume that TetA and TetR are part of the P sector of the proteome, which scales as 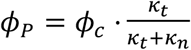. Without regulation of the synthesis rates *H*_*a*_ (*r*_*f*_) and *H*_*r*_ (*r*_*f*_), we would expect the concentrations of TetA and TetR to scale in the same way. Hence, we define 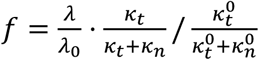 which leads to steady states proportional to 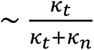 and simplifies to *f* = 1 at full nutrients and no drug. Example dynamics of these biochemical equations are shown in Supp. Fig. 6.

We combined the above biochemical equations with an agent-based model of growing and dividing cells. In this physical model, cells are represented by rod-shaped particles (a rectangle of varying length and width 2*R* with semi-circular caps of radius *R*) with diameter 2*R* = 1 *μm*. Volume exclusion is implemented by Hertzian repulsion forces ∼ (2*R* − *d*)^3/2^ for *d* ≤ 2*R*, where *d* is the distance between the centerlines (from cap center to cap center) of two cells. We consider the incompressible limit where repulsion is so strong that the overlap between cells is minimal at all times. Consequently, the resulting dynamics are insensitive to the exact force law for repulsion and mechanical parameters of the cells. Cells grow from a total length of 2 *μm* to 4 *μm* before they divide, with an elongation rate that leads to a doubling time of *ln*(2)/*λ* with *λ* from above. At birth, the nominal growth rate *λ*_0_ is chosen randomly within a window of ±25% for each cell in order to avoid synchronization of division, by multiplying the above-mentioned base values 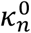 and 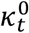 by a factor [0.75,1.25]. Cells are removed from the system when they reach the top of the domain. The left, right and bottom wall of the domain are modeled with Hertzian repulsion forces as well.

Each cell is equipped with the biochemical reaction scheme introduced above. Nutrients are tracked throughout the domain by a continuous field *c*(*x, y, t*) which follows the diffusion equation 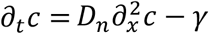 with a boundary condition *c* = 1 (full normalized nutrient concentration) at the top boundary of the domain. The local consumption rate *γ*(*x, y, t*) is determined by the state of the corresponding cells: Each cell consumes nutrients at a rate *γ*_0_ * *A* * *λ*/*λ*_0_, where *γ*_0_ is the nominal nutrient consumption rate at full nutrients and no drug (per unit time and unit area), and *A* and *λ* are the current area and growth rate of the cell, respectively.

An Euler method with adaptive time stepping was used for the evolution of the physical equations of motion of the cells and the biochemical reactions. The size of the time step was capped at a maximum of 10^−4^ min, and the maximum displacement of the cells was kept below 0.02μm by decreasing the time step as necessary. The diffusion equation for the continuous nutrient field was solved using a forward-time centered-space finite-difference scheme on a grid with a spatial discretization step size of 1μm in both directions and the same time step as for the equations of motion of the cells, using a standard 5-point stencil in two dimensions. The model was implemented in Julia (Bezanson et al. 2017).

Remaining model parameters, from (Schultz, Palmer, and Kishony 2017) where applicable: *K*_*i*_ = 0.015 *min*^−1^, *K*_*a*_ = 50 *min*^−1^, *k*_*a*_ = 10 μ*M,K*_*d*_ = 0.001 μ*M, K*_*ri*bo_ = 1 μ*M*, 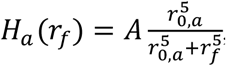, *A* = 0.0008 μ*M*/*min, r*_0,*a*_ = 0.0001 μ*M*, 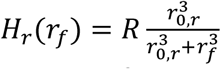, *R* = 0.0003 μ*Mmin*^−1^, *r*_0,*a*_ = 0.000075 μ*M, D*(*t*) = 0 *or* 50μ*M, D*_*n*_ = 1200 μ*m*^2^/*min, γ*_0_ = 0.4 *a. u*. μ*m*^−2^*min*^−1^, *c*_0_ = 0.01 *a. u*. μ*m*^−2^

## 6 Author Contributions

D.S. and P.B. designed the study. P.B. and J.H. designed the experimental technology, P.B. performed the experiments, M.S. and D.S analyzed the data. M.S., D.S. and P.B. designed the mathematical model, T.B.-L., L.H., and P.B. implemented the model and ran the simulations. M.S., D.S and P.B. wrote the manuscript.

## 8 Acknowledgments

The authors would like to thank Jonas Isensee for technical help with the combined simulations of agents and continuous field.

