## Supplementary Material for "Nutrient gradients mediate creation of highly resistant layers in structured microbial populations during antibiotic exposures"

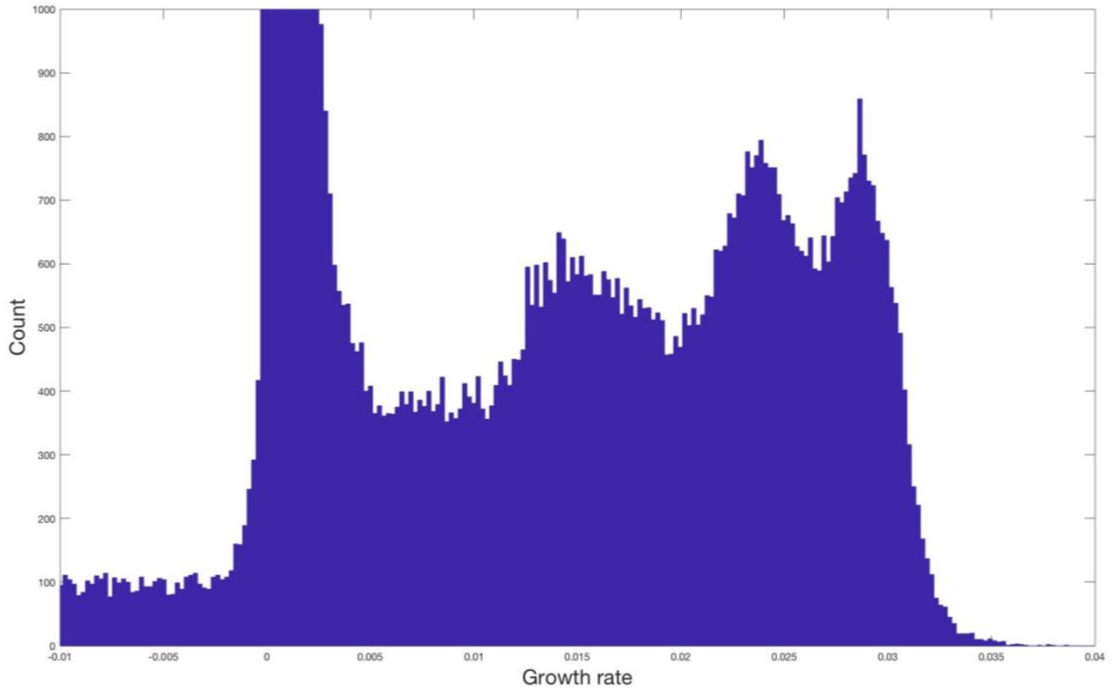


**Supplementary Figure 1. Histogram of growth rates across the whole trap, throughout the experiment.** A high peak at zero growth (cropped) corresponds to the arrested cells at the bottom of the trap. Peaks at 0.029 $min^{-1}$ and 0.023 $min^{-1}$ correspond to the steady-state growth rate of *tet* resistant *E. coli* in the absence and presence of 100µg/ml tetracycline, respectively.

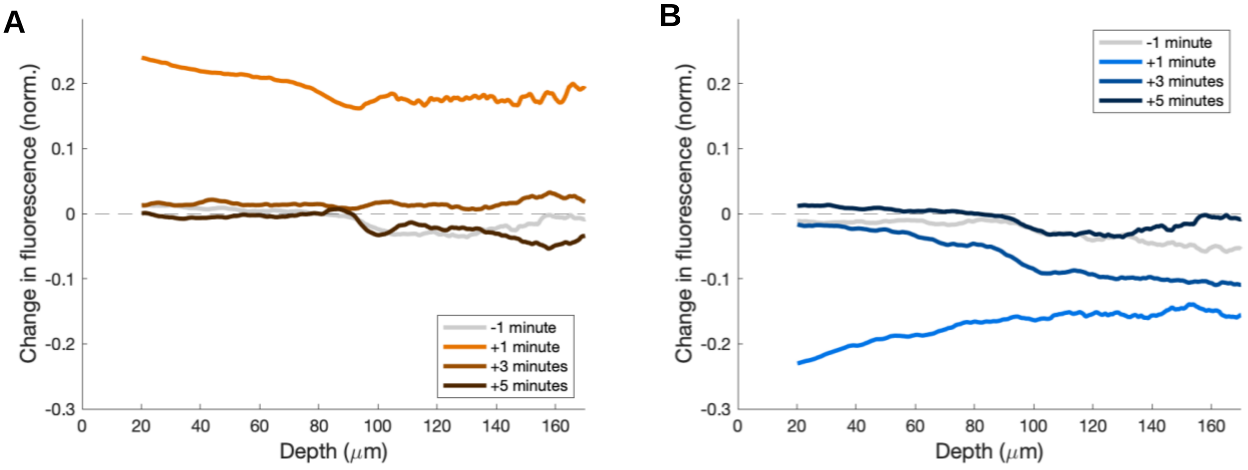


**Supplementary Figure 2. Timescale of the diffusion of small molecules into and out of the trap.** We measured changes in red fluorescence across the trap in the time points following (A) addition and (B) removal of sulforhodamine in the feeding channel, a dye with similar molecular weight as tetracycline. When the dye is added, it freely diffuses through the trap within the first minute, and no further changes in fluorescence are observed after the first measurement. When the dye is removed, it is mostly absent in the lower depths of the trap by the first measurement, but some dye remains in the back of the trap among dormant cells, likely due to boundary layer effects typical of laminar flow. This lingering dye then diffuses out by the second measurement, and no further changes are observed. Even during removal, the timescale of diffusion of small molecules out of the trap (~ 3 minutes) is still much smaller than the other relevant processes for the collective dynamics of the microcolony.

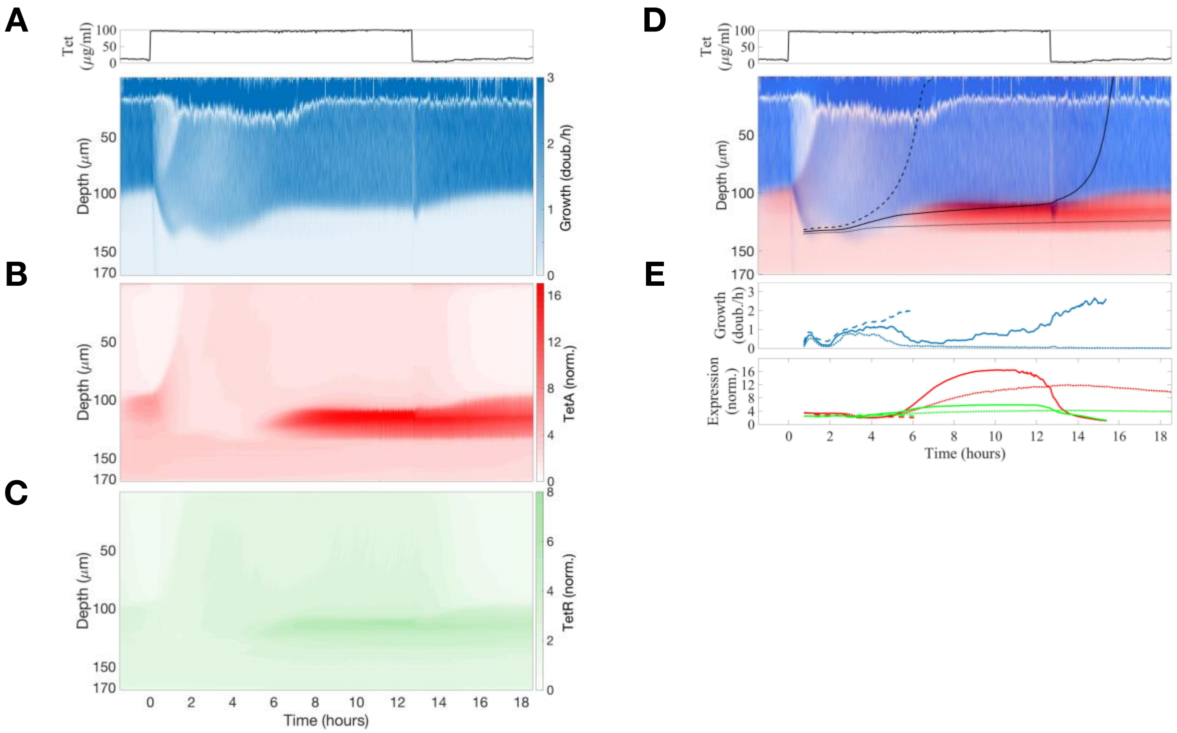


**Supplementary Figure 3. Repeat experiment in a second trap with a single long exposure**. Measurements of (A) cell growth, (B) TetA expression and (C) TetR expression during a single long exposure showed the same patterns depicted in Figure 2 in the main text. (D) Superimposed kymograph of cell growth (blue) and TetA expression (red), with black lines showing three approximate trajectories followed by single cells, calculated from the velocities. (E) Cell growth and expression of resistance genes TetR (green) and TetA (red) along these trajectories.

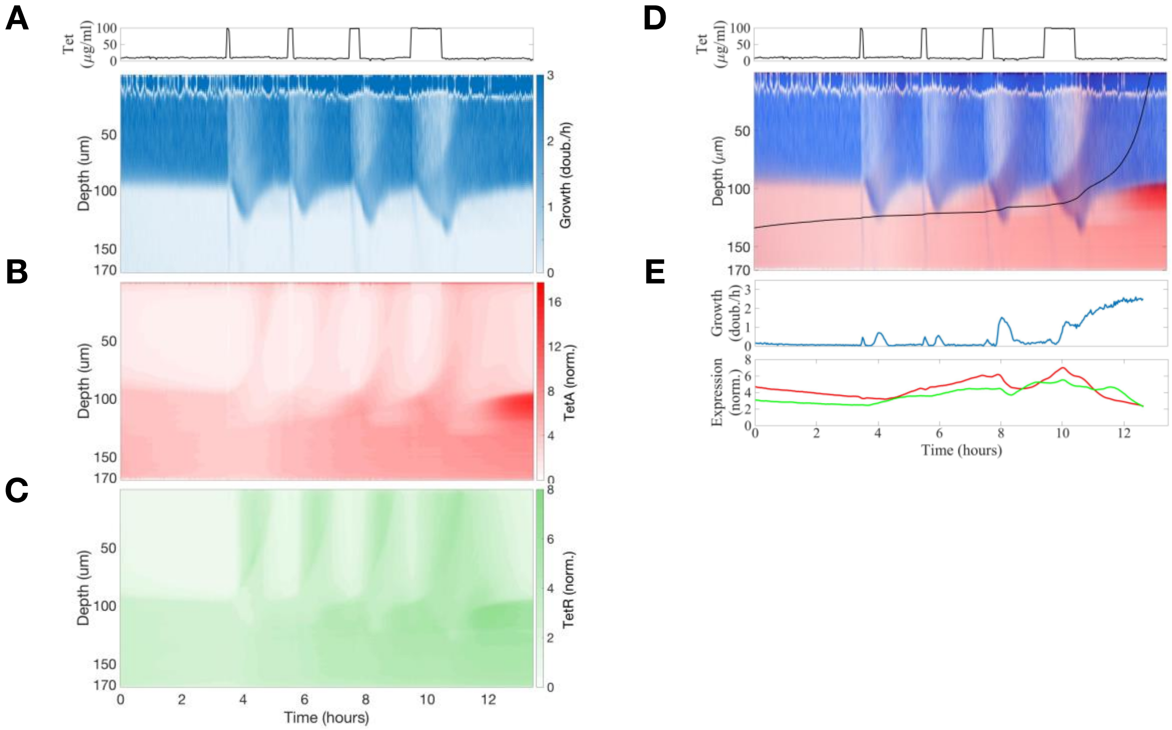


**Supplementary Figure 4. Repeat experiment in a second trap with pulses of tetracycline**. Measurements of (A) cell growth, (B) TetA expression and (C) TetR expression during a sequence of tetracycline exposures, with pulses of 5 min, 10 min, 20 min and 60 min delivered every 2 hours, showed the same patterns as Figure 4 in the main text. (D) Superimposed kymograph of cell growth (blue) and TetA expression (red), with a black line showing an approximate trajectory followed by a single cell originating in the interior of the colony, calculated from the velocities. (E) Cell growth and expression of resistance genes TetR (green) and TetA (red) along this trajectory.

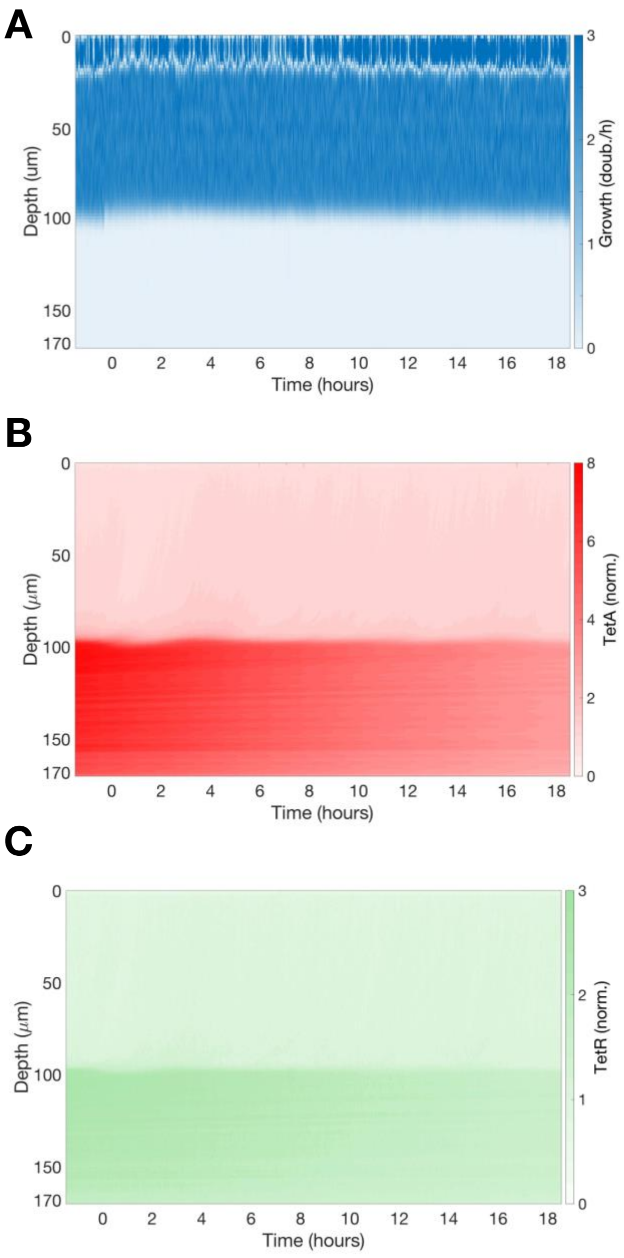


**Supplementary Figure 5. Control experiment showing a trap not exposed to tetracycline**. (A) A relatively sharp boundary between growing and arrested cells is maintained indefinitely in the absence of tetracycline. A slight reduction in both (B) mCherry and (C) GFP fluorescence measured in the bottom of the trap is presumably due to photobleaching, as arrested cells show reduced protein synthesis.

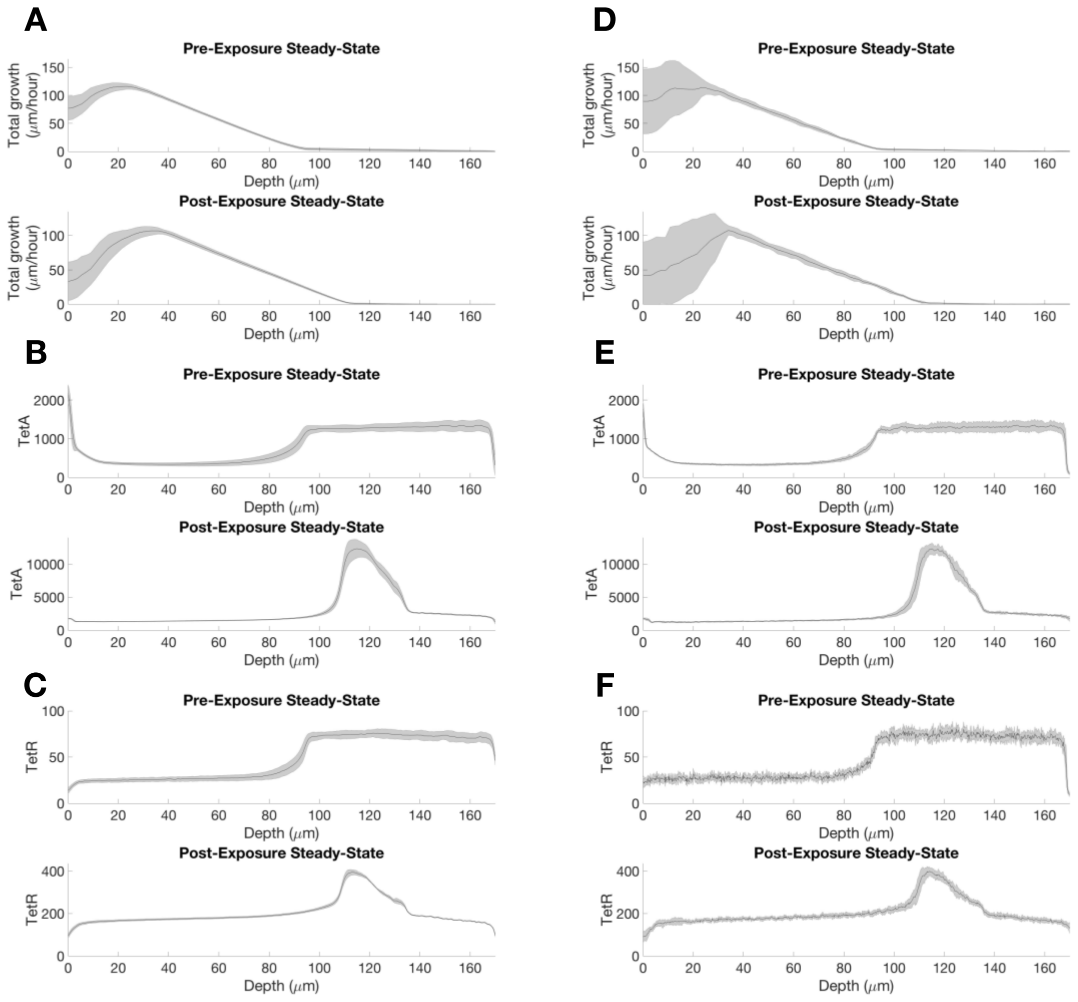


**Supplementary Figure 6. Variability in total growth, tetA and tetR measurements.** (A-C) Variability of measurements over time. We first averaged measurements for each depth across the width of each image. Then, using 104 images during a pre-exposure steady state, and 175 images during post-exposure steady state, we computed the mean and standard deviation of measured values over time at each depth, for total (A) cell growth (velocity), (B) TetA and (C) TetR. (D-F) Variability of measurements across a single image. We picked a representative image from both the pre-exposure steady state and the post-exposure steady state. Since there is little movement in the horizontal axis, we can segment the trap into 8 vertical slices and treat them as separate experiments. For each image and at each depth, 96 data points along the width of the image were available for total growth (velocity) and 460 for TetA and TetR fluorescence. We than calculated average values of each measurement at each depth within each slice, to smooth over the spatial structure of single cells, and used these 8 values to calculate the mean and standard deviation across the image.

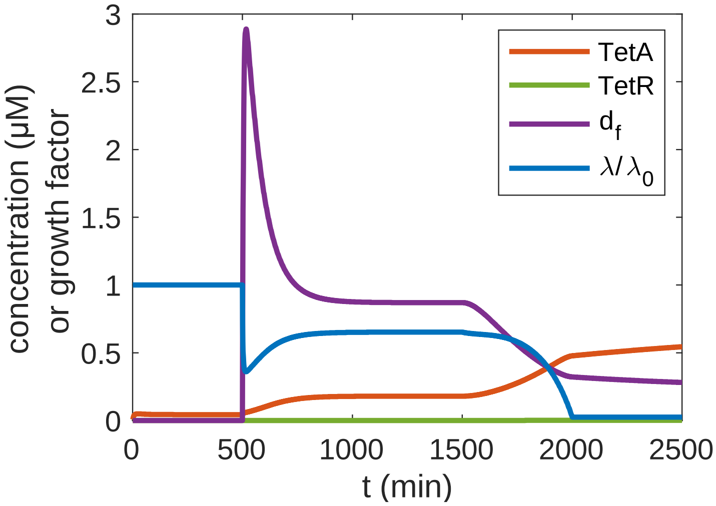


**Supplementary Figure 7. Example dynamics of the biochemical model equations**. At t=500, an indefinite tetracycline pulse is initiated. The growth rate drops transiently until enough TetA is expressed to decrease the intracellular drug concentration to its steady state level. From t=0 to t=1500, full nutrients are assumed, i.e. $\kappa_n = \kappa_n^0$. From t=1500 to t=2000, $\kappa_n$ is continuously decreased to 1% of its original level to demonstrate the effect of growth reduction through nutrients on expression in the presence of tetracycline.

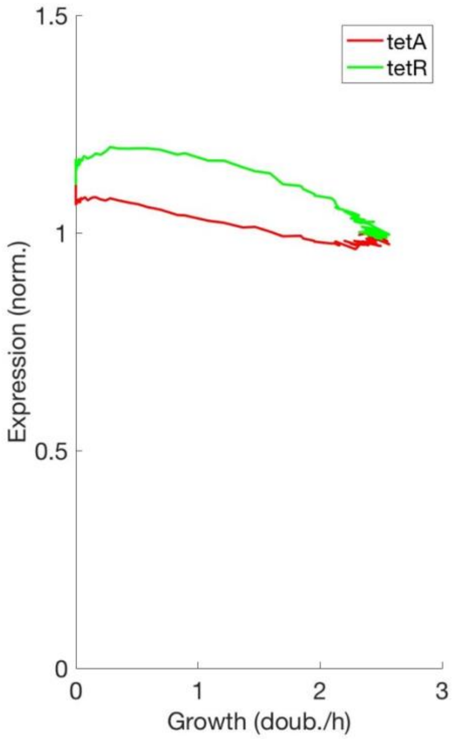


**Supplementary Figure 8. Steady-state expression of resistance reflects proteome partition.** Functional relationship between expression of TetA/TetR and cell growth, calculated from the model, obtained from the steady state before any contact with the drug. TetA and TetR expression decrease linearly with the cell growth rate, as the cells diverts more resources to the synthesis of ribosomes, according to the theory of proteome partition. Slightly increased expression values in our experimental system might reflect short exposures to the drug during the setup of the device.

**Supplementary Movie 1. Microfluidic device**. Microcolony of *tet* resistant *E. coli* cells growing in a trap in the microfluidic device. The colony was first subjected to windows of 5min, 10min, 20min and 60min of 100µg/ml tetracycline exposure, delivered every 2 hours, and then to a long exposure of 12 hours. The windows of tetracycline exposure are visible by the lack of red fluorescent dye in the media channel. Expression of efflux pump TetA is indicated in red, and repressor TetR is indicated in green.

**Supplementary Movie 2. Agent-based simulation of the microcolony**. Agent-based simulation showing transient reorganization by the growth pattern during a short drug pulse (t=5h) followed by a long pulse (t=8.5h) of tetracycline. Blue color indicates growth rate, and red color indicates expression of the efflux pump TetA. Cells tracked to yield the time traces in panels B-E in the main text are enclosed by a square of the same line style. The parameters used in this simulation are also described in the main text.
